# A single-dose intramuscular immunization of pigs with lipid-nanoparticle DNA vaccines based on the hemagglutinin antigen confers complete protection against challenge infection with the homologous influenza virus strain

**DOI:** 10.1101/2023.08.22.554288

**Authors:** The Nhu Nguyen, Sushmita Kumari, Sarah Sillman, Jayeshbhai Chaudhari, Danh Cong Lai, Hiep L.X. Vu

## Abstract

Influenza A virus of swine (IAV-S) is highly prevalent and causes significant economic losses to swine producers. Due to the highly variable and rapidly evolving nature of the virus, it is critical to develop a safe and versatile vaccine platform that allows frequent updates of the vaccine immunogens to cope with the emergence of new viral strains. The main objective of this study was to assess the feasibility of using lipid nanoparticles (LNPs) as a nanocarrier to deliver DNA plasmid encoding the viral hemagglutinin (HA) gene in pigs. Intramuscular administration of a single dose of the LNP-DNA vaccines resulted in robust systemic and mucosal in pigs. Importantly, the vaccinated pigs were fully protected against challenge infection with the homologous IAV-S strain, with only one out of 12 vaccinated pigs shedding a low amount of viral genomic RNA in its nasal cavity. No gross or microscopic lesions were observed in the lungs of the vaccinated pigs at necropsy. Thus, the LNP-DNA vaccines are highly effective in protecting pigs against the homologous IAV-S strain and can serve as a promising platform for the rapid development of IAV-S vaccines.

**Importance:** Influenza A virus of swine (IAV-S) is a significant pathogen of swine. The virus also poses a great public health concern due to its zoonotic potential. Although whole-inactivated virus (WIV) vaccines are available to control IAV-S, their heterologous efficacy is limited due to the substantial genetic/antigenic variation of the viral genome. This study provides compelling evidence demonstrating that lipid nanoparticle encapsulated DNA (LNP-DNA) vaccines induce robust systemic and mucosal immunity and complete protection of pigs against challenge infection with the homologous IAV-S strain. Importantly, the LNP-DNA vaccine approach meets the regulatory requirements to serve as an adaptable platform that can be frequently updated to match the emergence of new IAV-S variants. Thus, this LNP-DNA vaccine strategy exhibits great potential in effectively mitigating the economic impact of IAV-S on the swine industry and the associated public health threat.

## Introduction

Influenza A virus of swine (IAV-S) is an important respiratory pathogen that causes significant economic losses to swine producers and poses a great concern to public health due to its zoonotic potential (1). Three major types of IAV-S, H1N1, H1N2, and H3N2 are cocirculating in the swine population. The H1 subtype is further divided into three major lineages (1A, 1B, and 1C) with numerous genetic clades within each H1 lineage (2). Similarly, the H3 subtype is divided into multiple lineages (3). The profound genetic and antigenic diversity and the constant emergence of new variants pose a great challenge to the development of a broadly protective vaccine against IAV-S.

Whole-inactivated virus (WIV) vaccines are commonly used to control IAV-S (4). The WIV vaccines contain multiple IAV-S strains from different subtypes to enhance antigenic coverage. However, the WIV vaccines often fail to confer optimal levels of heterologous protection due to the substantial genetic and antigenic variation among IAV-S isolates circulating in the field (2, 5, 6). One approach to improve vaccine efficacy is frequently updating the vaccine immunogens to match the emergence of new viral strains. Due to the cost and the time demand of the vaccine licensing process, WIV vaccines are not updated in time to cope with the continual evolution of the viruses.

Recently, the United States Department of Agriculture’s Center for Veterinary Biologics (USDA-CVB) has issued two memoranda (Memorandum No. 800.213 and Memorandum No. 800.214, both signed on 03/12/2018) that provide guidance for licensing of non-replicating, nonviable biological veterinary vaccines. According to these memoranda, once a non-replicating, nonviable vaccine production platform has been established and the initial product has been licensed, licensure of new products containing sequence variants of the same vaccine antigens can be expedited, providing that there are no changes in the manufacturing process. These new regulations sparked interest in the development of novel non-replicating, nonviable platforms for veterinary vaccine development.

The DNA plasmid is noninfectious, nonviable, and safe to use in animals. However, vaccination of animals with the naked DNA plasmid often results in poor immune responses due to ineffective cellular uptake. Various methods have been developed, including gene guns or *in vivo* electroporation, to improve DNA plasmid delivery in swine (7–13). Recently, lipid nanoparticles (LNPs) have emerged as a promising nanocarrier for nucleic acid-based vaccines, particularly after the success of the COVID-19 vaccine (14, 15).

LNPs are versatile nanocarriers that have been utilized for the delivery of various nucleic acids, including siRNA, microRNA, mRNA, and DNA (14). These nanoparticles protect the encapsulated nucleic acids from degradation and enable their attachment and internalization into target cells, primarily through endocytosis. LNPs generally consist of four types of lipids: a cationic lipid, a phospholipid, cholesterol, and a polyethylene glycol (PEG)-conjugated lipid (16). Saturated phospholipids, such as distearoylphosphatidylcholine (DSPC), have high melting temperatures therefore they are used to construct highly stable liposomes and LNPs (17). Cholesterol stabilizes lipid bilayers by filling in gaps between phospholipids (17). Furthermore, cholesterol has the potential to crystallize on the surface of LNP, enhancing the endosomal escape of the nucleic acid cargo (18). PEG-lipids are used to enhance particle stability and prevent particle aggregation during preparation and storage, as well as to improve the circulation half-life and distribution of LNPs in vivo (16). Cationic lipids are positively charged, allowing them to interact electrostatically with the negatively charged phosphate backbone of nucleic acids and facilitate the incorporation of nucleic acids into the nanoparticles (16). There are two groups of cationic lipids: permanently ionized cationic lipids and conditionally ionized (ionizable) cationic lipids. The types and molar ratios of lipids used to formulate LNPs influence the uptake and endosomal releases of the encapsulated nucleic acid cargo (19). Research on LNP formulations for siRNA delivery has revealed that LNPs containing permanently ionized cationic lipids are more efficient for cellular uptake, while LNPs containing conditionally ionized cationic lipids are better for endosomal release. As a result, LNP-siRNA formulated by using a combination of permanently and conditionally ionized cationic lipids exhibits significantly improved gene silencing efficacy compared to LNP-siRNA based on a single type of cationic lipid (19).

In this study, we used a DNA plasmid encoding the hemagglutinin (HA) antigen of an IAV-S H3N2 strain as a model antigen to assess the effectiveness of two different LNP formulations, each with a unique combination of permanently and conditionally ionized cationic lipids. We demonstrated that a single-dose intramuscular administration of the LNP-DNA vaccine induced high titers of antibodies against the H3 antigen within 7-14 days post-vaccination. Furthermore, pigs vaccinated with the LNP-DNA vaccine were completely protected against challenge infection with the homologous H3N2 strains. These findings suggest that LNP-DNA can serve as an effective platform for the development of IAV-S vaccines.

## Results

### Generation and in vitro characterization of the lipid nanoparticle – DNA vaccines

We conducted an initial screening of multiple LNP-DNA formulations and identified two formulations that demonstrated transfection efficiency in HEK-293T cells. In LNP-DNA formulation 1 (LNP1), MC3, DOTAP, DSPC, cholesterol, and DMG-PEG2000 were mixed at a molar ratio of 35:5:10:48:2 (mol%), respectively while in LNP-DNA formulation 2 (LNP2), MC3, DOTAP, DSPC, cholesterol, DMG-PEG2000 were mixed at a molar ratio of 42:10:8:38:2, respectively.

The average diameter of LNP1 was 81 nm, which was slightly larger than LNP2 (76 nm) (Figure 1A). Both LNP1 and LNP2 formulations had a polydispersity index below 0.1, indicating a monodispersity distribution (Figure 1B). The zeta potentials of LNP1 and LNP2 were found to be within the neutral range, with mean values of -1.25 mV and +2.5 mV, respectively, in 100 mM Tris-Cl at pH 7.4 (Figure 1C). The mean encapsulation efficiencies of LNP1 and LNP2 were 62% and 82%, respectively (Figure 1D). Overall, LNP1 and LNP2 had similar physical characteristics.

**Figure 1:**
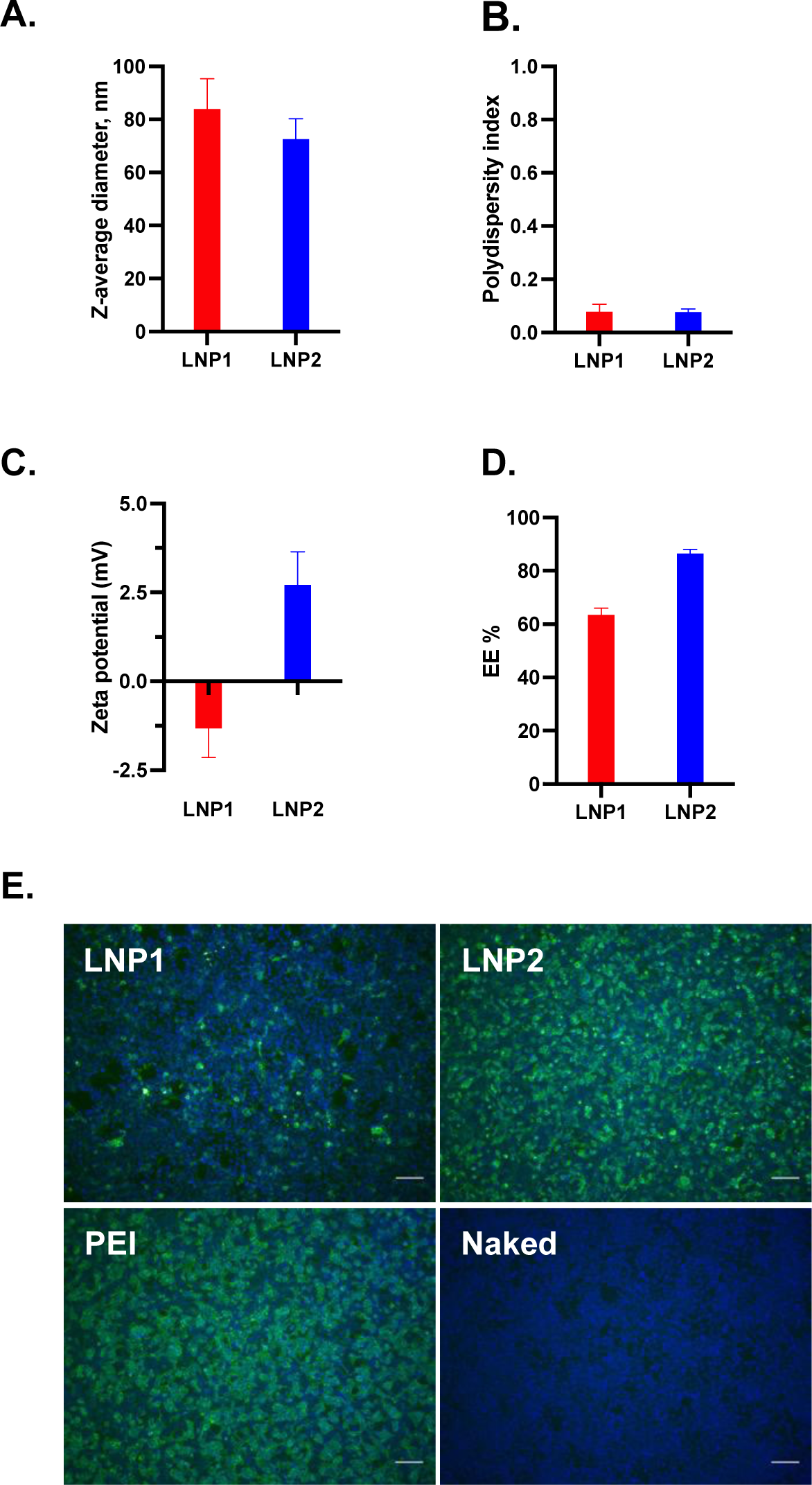
Physical characterization of the LNP-DNA vaccines and *in vitro* transfection efficiency. (A) Particle diameter determination by the nanoparticle tracking analysis (NTA). (B) Polydispersity index. (C) Zeta potential analysis. (D) Encapsulation efficiency (EE%). (E) Transfection efficiency in HEK-293T cells.

To evaluate the transfection efficiency, 0.5 μg encapsulated LNP1 or LNP2 vaccine was added directly into the medium of HEK-293T cells in a 24-well plate. Naked DNA plasmid was used as a negative control while DNA plasmid complexed with polyethyleneimine (PEI-DNA) was used as a positive control. At 48 hr post-transfection, the cells were fixed and stained with an anti-flag tag antibody to detect HA-expressing cells. As expected, no fluorescent positive cells were detected from cells transfected with naked DNA plasmid while approximately 90% positive cells were detected from cells treated with PEI-DNA (Figure 1E). The number of positive cells in LNP2-treated wells was slightly lower than in PEI-DNA transfected wells but was significantly greater than in LNP1-treated wells (Figure 1E). Thus, LNP2 had better *in vitro* transfection efficiency than LNP1.

### Pigs vaccinated with LNP-DNA vaccines elicited systemic and mucosal antibody responses

A vaccination/challenge experiment was conducted in 4-week-old pigs to assess the immunogenicity and protective efficacy of the LNP1 and LNP2 vaccines. We first measured HA-specific IgG in plasma samples using an indirect ELISA. As expected, anti-HA IgG antibodies were not detected in the PBS-control group at any sampling dates (Figure 2A). On the other hand, anti-HA IgG antibodies were detected in both LNP1 and LNP2 vaccinated pigs on day 7 post-vaccination and the antibody titers sharply increased, reaching the titers of 1:10^6^ on day 14 post-vaccination, and maintained at a similar titer until the end of the 35-day observation period. There were no significant differences in kinetics and magnitude of anti-HA IgG antibody responses between the LNP1- and LNP2-groups.

**Figure 2:**
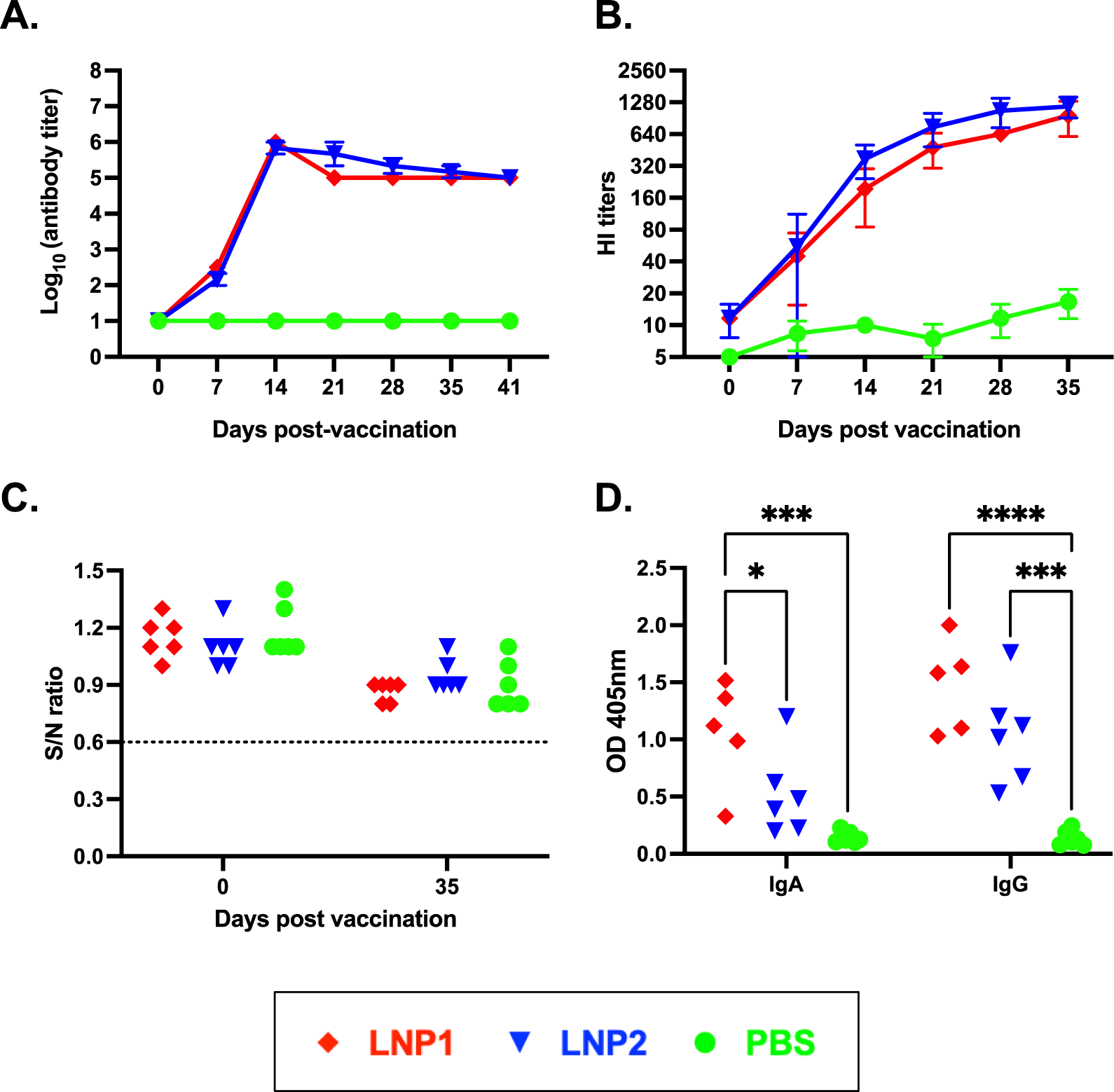
Systemic and mucosal antibody responses following vaccination. (A) Anti-HA IgG titers in blood samples collected on various days post-vaccination. Data are expressed as the log_10_ of the reciprocal of the highest plasma dilution at which anti-HA antibodies were observed. The samples were initially diluted at 1:100 and samples with undetectable antibodies at this dilution were considered negative and assigned a value of 1 log_10_. (B) Hemagglutinin inhibition (HI) antibody titers measured against the H3N2 TX98 virus. Data are expressed as the reciprocal of the highest plasma dilution at which hemagglutination inhibition was observed. The samples were initially diluted at 1:10 and samples with undetectable HI activity at this dilution were considered negative and assigned a value of 5. (C) Anti-NP antibody levels measured using a commercial ELISA kit. Data are presented as the sample to negative (S/N) ratio. The horizontal dotted line at S/N of 0.6 is the assay cutoff. Samples with S/N greater than 0.6 were considered to be negative. (D) HA-specific IgA and IgG in BALF collected at necropsy.

Next, we measure plasma HI antibody titers against the homologous H3N2 TX98 virus. Plasma HI antibodies were initially detected on day 7 post-vaccination and exhibited a gradual increase, reaching mean titers above 1:640 by day 35 post-vaccination (Figure 2B). No significant difference in HI titers was observed between the LNP1- and LNP2-groups. The PBS-control group showed low HI titers (1:10), potentially due to non-specific inhibition.

We also measured antibodies against the IAV-S nucleoprotein (NP) using a commercially available ELISA kit that is commonly used for serodiagnosis. All plasma samples collected before and on day 35 post-vaccination showed negative interpretations for the NP antibodies (Figure 2C). The results demonstrate that the pigs were not exposed to IAV-S and that the anti-HA IgG or HI antibodies detected in the plasma of pigs in the LNP1 and LNP2 groups can be attributed solely to the vaccination.

To assess the mucosal antibody response elicited by the LNP-DNA vaccines, we measured HA-specific IgA and IgG antibodies in BALF samples collected at necropsy using an indirect ELISA. Both HA-specific IgG and IgA antibodies were detected in BALF samples collected from LNP1-vaccinated pigs while only HA-specific IgG antibodies were detected in BALF samples of the LNP2-vaccinated pigs (Figure 2D). Collectively, the data demonstrate that both LNP1 and LNP2 vaccines elicited robust systemic antibody responses in the vaccinated animals. However, the LNP1 vaccine appeared to induce a better mucosal IgA antibody response than the LNP2 vaccine.

### Pigs vaccinated with LNP-DNA vaccines elicited strong T-cell responses

The frequencies of IFN-γ secreting cells in PBMCs were evaluated using the IFN-γ ELISPOT assay. No IFN-γ spots were detected in PBMCs collected before vaccination or in PBMCs collected from the PBS group on day 35 post-vaccination (Figure 3). The number of IFN-γ spots in PBMCs collected from the LNP1 or LNP2 groups on day 35 post-vaccination ranged between 50 and 350 spots per 10^6^ PBMCs, with no significant difference observed between these two groups.

**Figure 3:**
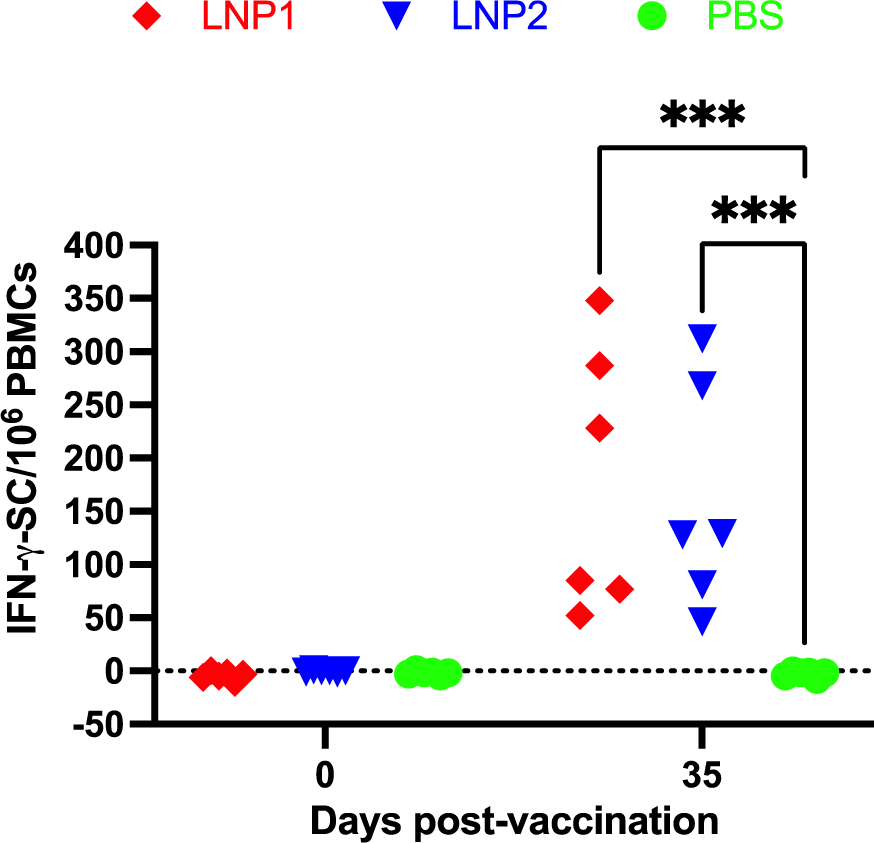
Virus-specific IFN-γ-secreting cell responses following vaccination. Data are expressed as the IFN-γ secreting cells (IFN-γ-SC) per 10^6^ PBMC cells. Data were analyzed by one-way ANOVA, followed by Tukey’s multiple comparisons test.

To further investigate the characteristics of the T-cell responses, multicolor flow cytometry was utilized to analyze various T-cell populations in the PBMCs, including CD4^+^, CD8+, and CD4^+^CD8^+^ double-positive, for their expression of the three important T-cell cytokines: IFN-γ, TNF-α, and perforin. Overall, cells expressing IFN-γ were mainly observed in CD4^+^ and CD8^+^, but not in CD4^+^CD8^+^ T-cell populations (Figure 4A&B). On the other hand, cells expressing TNF-α and perforin were only detected in CD8^+^ and CD4^+^CD8^+^, but not in the CD4^+^ T-cell population. When compared to the PBS group, significantly higher frequencies of IFN-γ-expressing cells were observed in both CD4^+^ and CD8^+^ T cells of the LNP1 group, as well as in CD4^+^ T cells of the LNP2 group (Figure 4 A&B). Both CD8^+^ and CD4^+^CD8^+^ T cells of the LNP1 and LNP2 groups displayed a higher frequency of TNF-α-expressing cells than the corresponding cell populations of the PBS group (Figure 4 E&F). Regarding perforin expression, only the CD8^+^ and CD4^+^CD8^+^ T-cell populations of the LNP1 group exhibited a greater frequency of perforin-expression cells than the PBS group (Figure 4 H&I). When comparing the LNP1 and LNP2 groups, statistically significant differences were only observed in the population of CD4^+^CD8^+^ T-cells expressing TNF-α and perforin (Figure 4 F&I).

**Figure 4.**
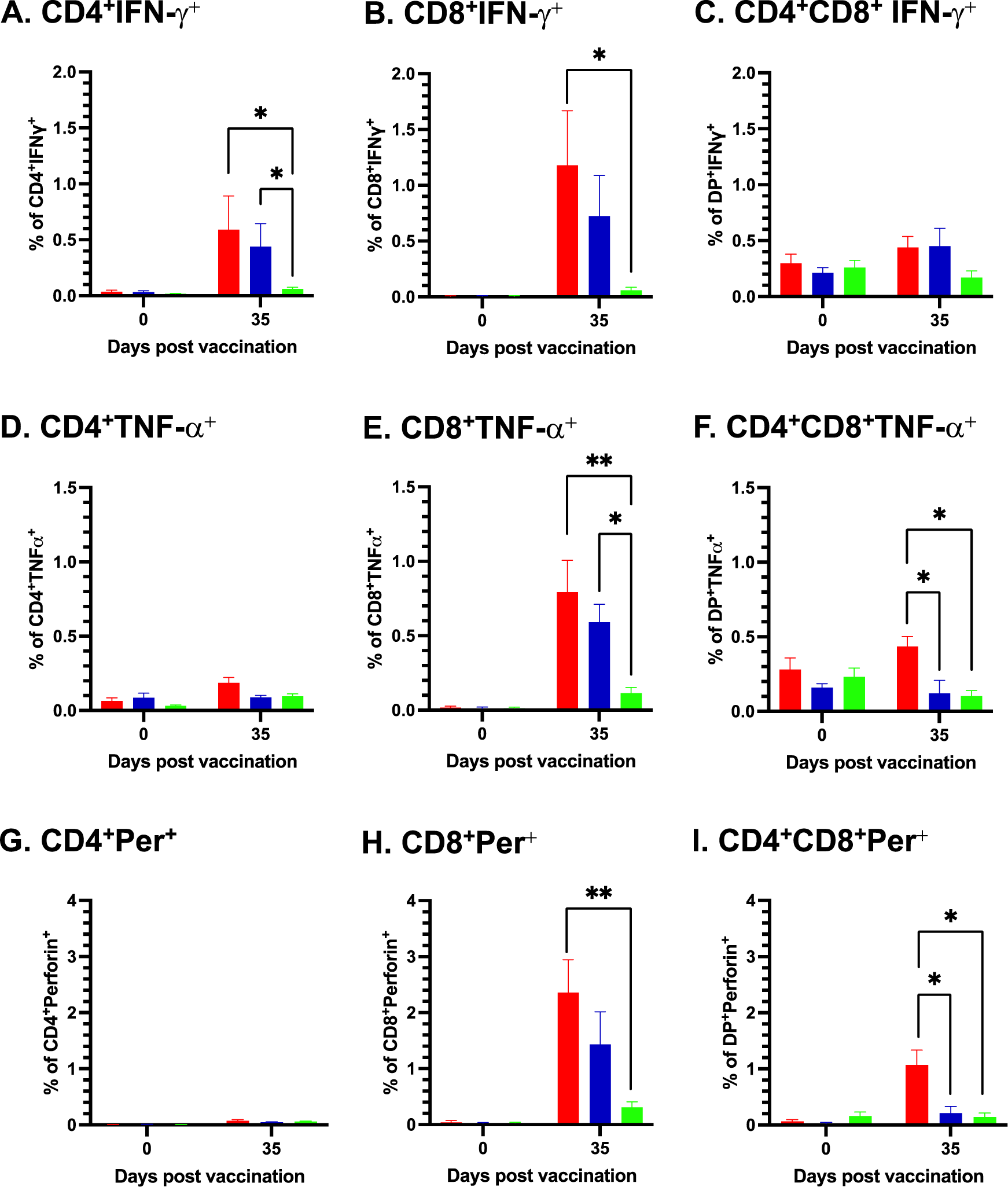
Characterization of the T-cell responses. PBMCs were stimulated with the H3N2 TX98 virus at an M.O.I of 2. The cells were then stained with antibodies against surface markers (CD3ε, CD4α, and CD8α), followed by staining with antibodies against three intracellular cytokines (IFN-γ, TNF-α, and perforin). (A-C). Frequency of cells expressing IFN-α, (D-F) Frequency of cells expressing TNF-α. (G-I) Frequency of cells expressing perforin (Per). Data are presented as means and the standard error of the mean of six animals in each group. Data were analyzed by the Kruskal-Wallis test, followed by Dunn’s multiple comparisons test. * *p* ≤ 0.05,** *p* ≤ 0.01.

### Pigs vaccinated with LNP-DNA vaccines were protected against challenge infection with the homologous influenza strain

On day 35 post-vaccination, all pigs were challenged with an intranasal/intratracheal inoculation with the homologous IAV-S strain H3N2 TX98. Daily nasal swabs were collected to measure viral shedding. Viral RNA was not detected in any samples collected before the challenge infection (Figure 5A). In the PBS group, viral RNA was detected from all samples starting from day 1 post-challenge, reached a maximal titer of 10^8^ copies/100 μL on day 3 post-challenge, and declined to 10^6^ copies/100 μL on day 5 post-challenge. In the LNP1 group, only one pig had a low copy number of viral RNA in samples collected on days 3 and 4 post-challenge. None of the pigs in the LNP2 group had detectable levels of viral RNA at any sampling dates (Figure 5A). Area-under-the-curve (AUC) of the nasal viral shedding was calculated for individual pigs during the course of five days post-challenge. The AUC of the LNP1 and LNP2 groups was significantly lower than that of the PBS group (Figure 5B).

**Figure 5:**
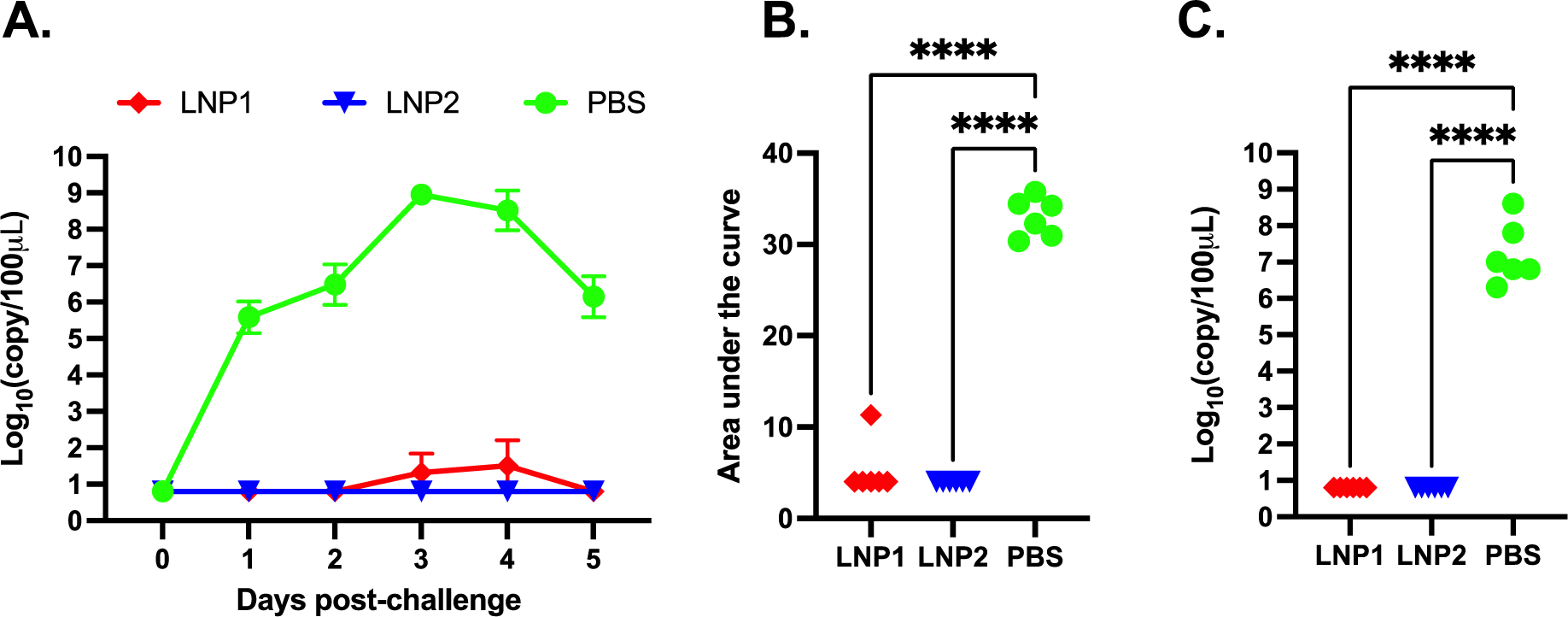
Viral shedding after challenge infection with H3N2 TX98. (A) Viral RNA in nasal swabs as determined by RT-PCR. Data are presented as log_10_ viral RNA copies per 100 μL of the sample. (B) Area under the curve of the nasal viral loads in each pig during the five days post-challenge infection with H3N2 TX98. (C) Viral RNA in bronchoalveolar lavage fluid (BALF) collected on day 5 post-challenge infection. For graphical and statistical purposes, samples with undetectable levels of viral RNA were assigned a value of 0.8 log_10_ which is equivalent to the assay detection limit.

On day 5 post-challenge, the pigs were humanely euthanized and necropsied. BALF samples were collected to evaluate viral loads within the lungs. High copy numbers of viral RNA, ranging from 10^6^ to 10^8^ copies/100 μL of BALF, were detected in all pigs from the PBS group. In contrast, viral RNA was not detected in any pigs from the LNP1 or LNP2 groups (Figure 5C).

At necropsy, gross lung lesion was scored by a board-certified pathologist blinded to the experimental design. Lungs of pigs in the PBS group presented with purple-red consolidation typical of IAV-S infection, with the percentage of total lung surface consolidation ranging from 0.4% to 4.58%. The consolidation was more prominent on the apical and middle lobes (Figure 6A). In contrast, only one pig in the LNP2 group had lung consolidation (2.49%), while none of the pigs from the LNP1 group had visible lung consolidation. The percentage of lung consolidation was not statistically different between the LNP1 and LNP2 groups, and both groups had significantly lower lung consolidation than the PBS group (Figure 6A).

**Figure 6:**
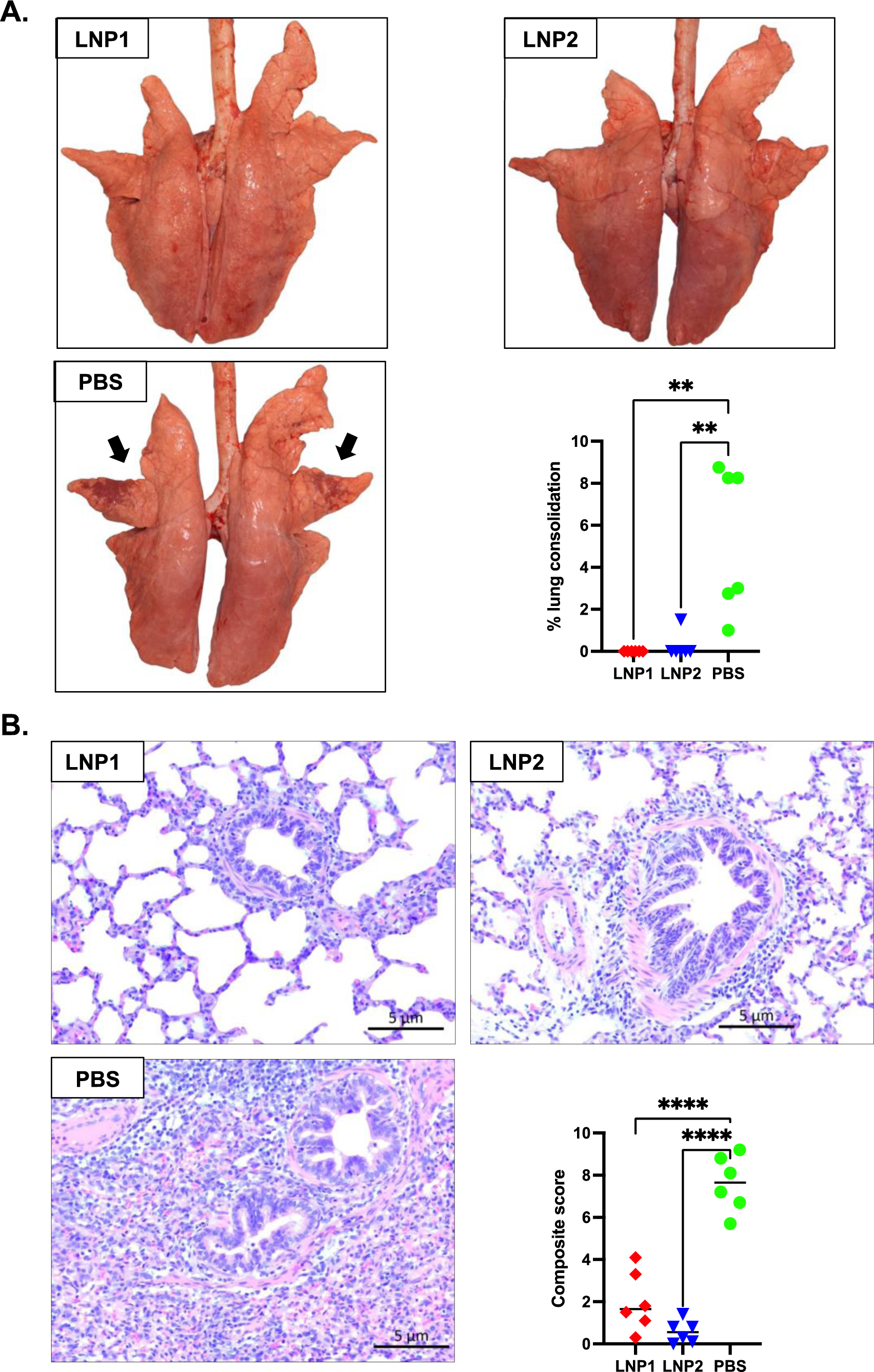
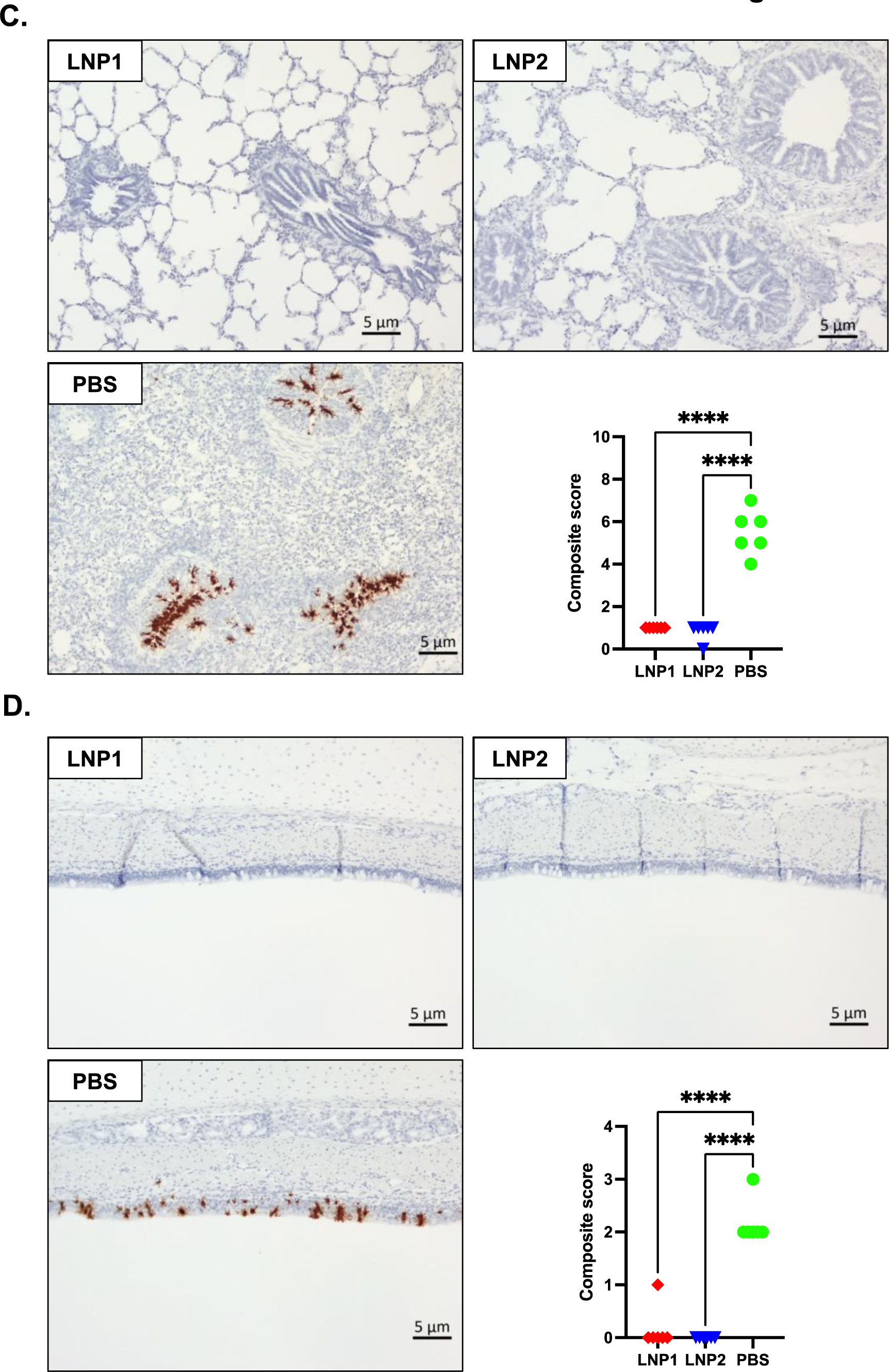
Lung pathology and presence of virus-infected cells in tissue samples. (A) Representative photos of lungs taken during necropsy. Black arrows indicate areas of the lungs with typical consolidation caused by IAV-S. The graph indicates the percentage of lung consolidation calculated based on the weighted proportions of each lobe to the total lung volume. (B) Representative images of lung sections stained with H&E and the composite microscopic lesion scores based on parameters described in the Materials and Methods. (C) Representative images of lung sections stained with ISH for the detection of viral NP mRNA transcript and the composite ISH scores. (D) Representative images of tracheal sections stained with ISH for the detection of viral NP mRNA transcript and the composite ISH scores.

Sections of three lung lobes (apical, middle, and caudal) from each pig were stained with H&E to evaluate microscopic changes. Lung sections from pigs in the PBS group exhibited variable, but typically moderate peribronchiolar lymphocytic cuffing with interstitial pneumonia, necrosis, and attenuation of epithelial cells in bronchioles, and areas of suppurative bronchiolitis. The multifocal interstitial pneumonia and consolidation were characterized by the infiltration of macrophages, lymphocytes, and neutrophils in the alveolar septae, sometimes spilling out into the alveolar lumens (Figure 6B). Conversely, lung sections of pigs from the LNP1 and LNP2 groups showed only mild interstitial pneumonia and peribronchial lymphocytic infiltration. The composite microscopic scores of pigs from the LNP1 and LNP2 groups were statistically similar and significantly lower than those of the PBS group (Figure 6B).

The *in-situ* hybridization (ISH) assay was used to detect virus-infected cells in the trachea and the middle lung lobe. Numerous virus-infected cells were detected in the tracheal, bronchial, and bronchiolar epithelium of pigs from the PBS group (Figure 6 C&D). On the other hand, virus-infected cells were rarely detected in either tracheal or lung sections of pigs from the LNP1 and LNP2 groups (Figure 6 C&D).

Collectively, the data clearly indicate that vaccination with either the LNP1 or LNP2 vaccines fully protected pigs against challenge infection with the homologous IAV-S strain.

## Discussion

There are several publications describing the immunogenicity and protective efficacy of DNA vaccines encoding the HA antigen in pigs. In several studies, naked DNA plasmids were administered intramuscularly or intradermally using a needle-free intradermal applicator (7–9, 20). In some instances, the DNA plasmids were adsorbed onto the surface of gold microparticles for delivery into the pig’s epidermis using a gene gun (11, 12). Electroporation has also been explored to enhance the DNA plasmid uptake (9). Additionally, the coding sequence of the vaccine immunogens was fused in-frame into flagellin or co-administered with a DNA plasmid expressing porcine interleukin 6 (pIL6) to enhance the immunogenicity of the DNA vaccines (12, 13). Up to 4 mg of DNA plasmid have been administered to pigs (20). Most of the time, HI antibody titers were not detected in DNA-vaccinated pigs after one immunization. Moreover, pigs vaccinated with the DNA vaccines were not protected from being infected with IAV-S after challenge infection although they may have shorter durations and lower magnitudes of virus shedding in their nasal cavity. It was reported in one study that viral RNA was not detected in the nasal swabs collected from pigs vaccinated with the DNA vaccine (7). However, in that particular study, viral RNA was not detected in nasal swabs collected from three out of five control pigs that were challenged with the IAV-S strain.

In this study, we assessed the effectiveness of two different LNP formulations as a nanocarrier to deliver the DNA plasmid encoding the HA antigen of the H3N2 TX98 in pigs. These two LNP formulations differed mainly in the molar ratios of permanently ionized cationic (DOTAP) and conditionally ionized cationic (MC3) lipids. Specifically, the LNP1 had a molar ratio of MC3 to DOTAP of 35:5 (mol%), and these two cationic lipids made up 40% (mol%) of the total lipid component. On the other hand, the LNP2 had a molar ratio of MC3 to DOTAP of 42:10 (mol%), making up 52% of the total lipid components. Accordingly, the molar ratios of two helper lipids, DSPC, and cholesterol, were adjusted to accommodate the difference in the percentage of cationic lipids used in the LNP1 and LNP2. Despite the differences in ratios of the lipid components, the physical characteristics of these two LNP products were very similar. However, LNP2 had significantly greater *in vitro* transfection efficiency than LNP1. Interestingly, when administered intramuscularly to pigs, the LNP1 and LNP2 vaccines elicited similar kinetics and magnitude of antibody and IFN-γ ELISPOT responses. Notably, pigs vaccinated with a single dose of either LNP1 or LNP2 vaccines mounted high HI antibody titers on day 14 post-vaccination and were fully protected against challenge infection with the homologous IAV-S strain, with only one out of 12 vaccinated pigs exhibiting a low copy number of viral RNA, ranging from 10^3.9^ to 10^5^ copies/100 μL of nasal swabs, detected on day 3 and 4 post-challenge, respectively. Our results demonstrate that the two LNP-DNA vaccines tested in this study induced complete protective immunity in pigs. The close similarity in the protective efficacy of the LNP1 or LNP2 vaccines suggests that, under the conditions of this study, *in vitro* transfection efficiency does not correlate with the protective potency of the LNP vaccines in pigs.

We utilized multicolor flow cytometry to analyze various T-cell populations in PBMCs, including CD4^+^, CD8^+^, and CD4+CD8^+^ cells, for their expression of three cytokines: IFN-γ, TNF-α, and perforin. IFN-γ is commonly used as a marker of T-cell responses in pigs, and TNF-α can augment the killing capacity of IFN-γ (21). Perforin is a marker of cytotoxicity T cells. Antigen-specific T-cells co-producing IFN-γ and TNF-α have been shown to be crucial for vaccine-induced protection against intracellular pathogens (22). Although we simultaneously stained the cells with antibodies against the three cytokines in this study, we did not detect a significant frequency of cells simultaneously expressing two or three cytokines because the frequencies of cells expressing each cytokine were relatively low. Overall, the results showed that pigs vaccinated with LNP1 exhibited statistically higher frequencies of cells expressing IFN-γ, TNF-α, and perforin than those in the PBS group. On the other hand, pigs vaccinated with LNP2 showed no significant difference in the frequencies of cells expressing these cytokines except for CD4+ T cells expressing IFN-γ and CD8+ T cells expressing TNF-α, which were significantly higher than the corresponding cell populations in the PBS group. Although pigs in the LNP1 and LNP2 groups exhibited similar numbers of IFN-γ-secreting cells, as measured by the ELISpot assay, pigs in the LNP1 group had higher frequencies of CD4^+^CD8^+^ T cells secreting TNF-α and perforin than those vaccinated with the LNP2 vaccine. Thus, the LNP1 vaccine may elicit a different T-cell response compared to the LNP2 vaccine, but it remains unclear how much such a difference in the T-cell response contributes to the protection of pigs against challenge infection.

In summary, we report here the development of two LNP-DNA vaccine formulations that elicit a robust antibody and T-cell response and completely protect pigs against challenge infection with the homologous IAV-S strain. Additional studies are being conducted to determine the minimal dose of LNP-DNA vaccine that is required to induce protection and the duration of the protective immunity.

## Materials and methods

### Cells, viruses and lipids

HEK-293T (ATCC CRL-3216) cells were used for the evaluation of transfection efficiency. Madin-Darby Canine Kidney (MDCK) cells (ATCC® CCL-34) were used for propagation and titration of IAV-S. The IAV-S A/swine/Texas/4199-2/1998 (H3N2 TX98) was obtained from the National Veterinary Services Laboratories (NVSL, Ames, IA). The virus was propagated and titrated in MDCK cells.

The lipids used in this study included DLin-MC3-DMA (MC3) (Nanosoft Polymer, Winston-Salem, NC), 1,2-dioleoyl-3-trimethylammonium-propane (DOTAP) (Cayman Chemical, Ann Arbor, MI), Cholesterol (Sigma Aldrich), distearoylphosphatidylcholine (DSPC), and 1,2-dimyristoyl-rac-glycero-3-methoxypolyethylene glycol-2000 (DMG-PEG2000) (Avanti Polar Lipids, Birmingham, AL). The lipids were separately dissolved in absolute ethanol.

### DNA plasmid construction

The H3N2 TX98 HA gene coding sequence (GenBank accession no. AEK70342.1) was codon optimized for optimal expression in swine cells (*Sus scrofa*). A flag-epitope sequence (DYKDDDDK) was fused in-frame to the 3’ end of the gene to facilitate protein detection. The gene fragment was chemically synthesized using a commercial DNA synthesis service (GenScript, Piscataway, NJ) and cloned into the pCI plasmid (Promega, Madison, WI). Large-scale DNA plasmid was amplified in *Escherichia coli* DH5α and purified by using a plasmid giga prep kit (Zymo Research, Costa Mesa, CA). DNA sequencing was performed to confirm the plasmid sequence’s authenticity.

### Preparation of LNPs

Two LNP-DNA formulations were prepared following a method described previously with some modifications (23). In formula 1 (LNP1), specific volumes of MC3, DOTAP, DSPC, cholesterol, and DMG-PEG2000 were mixed at a molar ratio of 35:5:10:48:2 to form a final mixture with a total lipid concentration of 28 mM. In formula 2 (LNP2), MC3, DOTAP, DSPC, cholesterol, and DMG-PEG2000 were mixed at a molar ratio of 42:10:8:38:2 to form a final mixture with a total lipid concentration of 25 mM. DNA plasmid encoding HA antigen of the H3N2 TX98 was diluted in 50 mM citrate buffer pH 4.0 for LNP1 or in 25 mM sodium acetate buffer pH 4.0 for LNP2. The lipids (organic phase) and DNA solution (aqueous phase) were mixed by using a mixer-4 chip and the NanoGenerator™ Flex-M Nanoparticle Synthesis System (Precigenome, San Jose, CA). The DNA and lipid flow rates were set at 3:1 (v/v), and the total flow rate was set at 4 mL per minute. The nitrogen-to-phosphate (N/P) ratio (mol/mol) was fixed at 4.5. The resulting products were dialyzed against 100 mM Tris-Cl buffer at pH 7.4 (LNP1) or 100 mM Tris-Cl buffer containing 10% sucrose at pH 7.4 (LNP2) using the Slide-A-Lyzer^TM^ G2 dialysis cassettes with the molecular weight cut-off of 10 kDa (Thermo Fisher Scientific, Carlsbad, CA). After dialysis, LNPs were passed through 0.45 μm PES filters and the encapsulated DNA plasmid concentration was adjusted to 100 μg/mL.

### LNP characterization

LNP sizes were measured by following the nanoparticle tracking analysis (NTA) method by using the Flow NanoAnalyzer (NanoFCM, Tokyo, Japan). The polydispersity index (PDI) and zeta potential (mV) were determined using the Malvern Zetasizer^®^ (Nano ZS, Malvern Instrument, Worcestershire, UK). DNA plasmid encapsulation efficiency (EE%) was quantified using a Quant-iT™ PicoGreen® dsDNA assay kit (Thermo Fisher Scientific) as described previously (23).

### Transfection efficiency in vitro

To assess the transfection efficiency of the LNP1 and LNP2 vaccines, 500 ng of encapsulated DNA was directly added to one well of the 24-well plate containing HEK-293T cells. An equal amount of naked-DNA plasmid was added to another well to serve as a negative control while DNA plasmid mixed with PEI transfectant (PEI-DNA) was used as a positive control. At 48 hr post-transfection, the cells were washed with phosphate-buffered saline (PBS) and fixed with a mixture of methanol and acetone (1:1 v/v), and subjected to an indirect immunofluorescence assay (IFA) using an anti-flag tag antibody and the Alexa fluor^TM^ 488 labeled goat anti-mouse IgG (H+L). The cell nucleus was stained with DAPI (4′,6′-diamidino-2-phenylindole). Fluorescence images were captured separately using a green or a blue filter and then merged to create the final composite images.

### Animal experiment

Eighteen weaned pigs, approximately 4 weeks old and seronegative for porcine reproductive and respiratory syndrome virus (PRRSV) and IAV-S, were obtained from Midwest Swine Research and housed in the animal biosafety level 2 (ABSL2) research facility at the University of Nebraska-Lincoln (UNL). The pigs were randomly assigned into three groups of six pigs. Pigs in groups 1 and 2 were intramuscularly injected with the LNP1 or LNP2 vaccines, respectively. Each dose of the LNP vaccine contained 500 µg of encapsulated DNA. Group 3 was injected with PBS to serve as a non-vaccination control.

Whole-blood samples were collected from all pigs before and weekly after immunization using vacuum tubes containing ethylenediaminetetraacetic acid (EDTA) anticoagulant. The blood tubes were then centrifuged at 1200 × g for 15 minutes, and the plasma was collected and stored at -20°C for evaluation of humoral immune responses. The cells were resuspended in PBS and peripheral blood mononuclear cells (PBMCs) were collected and washed twice with PBS containing 2% FBS as described before (24). The cells were then suspended in a cell freezing medium containing 50% RPMI, 40% FBS, and 10% DMSO, and cryopreserved for evaluation of T cell responses.

On day 35 post-vaccination, all pigs were challenged by a combination of intratracheal and intranasal inoculation of 2 × 10^5^ TCID_50_ of the H3N2 TX98 virus diluted in 4 mL serum-free DMEM. To administer the challenge inoculation, the pigs were sedated by intramuscular injection with telazol, ketamine, and xylazine. Two mL of the virus inoculum were administered via an endotracheal tube inserted into the tracheal tract, and 1 mL of the inoculum was administered into each nostril.

Nasal swabs were taken from all pigs daily post-challenge to measure viral shedding. On day 5 post-challenge, the pigs were humanely euthanized with an overdose of sodium pentobarbital. The lungs were removed, and bronchioalveolar lavage fluid (BALF) samples were collected using 50 mL of cold PBS to measure viral load in the lungs. Gross lung lesion was visually estimated and samples of the trachea (approximately 1 inch above the carina), left apical, middle, and caudal lung lobes were collected and fixed in 10% buffered formalin for histopathologic examination.

### Pathological analysis

The evaluation of pathology parameters was conducted by a veterinary pathologist, who serves as a coauthor but was intentionally kept blinded to the identity of the experimental groups during the scoring process. For the evaluation of gross lung lesions, the percentage of purple-red consolidation typical of IAV-S infection was visually estimated for each lung lobe. The total percentage of lung surface affected was then calculated based on the weighted proportions of each lobe to the total lung volume (25).

For the evaluation of microscopic lung lesions, sections of the apical, middle, and caudal lung lobes were stained with hematoxylin and eosin (H&E) using routine procedures. Each section was scored for six parameters, including peribronchiolar lymphocytic cuffing, interstitial pneumonia, bronchial and bronchiolar epithelial cell changes, bronchiolitis, edema, and epithelial exocytosis, following the scoring scale described previously (26). The average composite scores of the three lobes are reported.

Virus-infected cells were detected in lung and trachea sections using the RNA *in situ* hybridization (ISH) assay as previously described (27). The frequency of virus-infected cells in airway epithelium and pulmonary parenchyma was estimated for each lung component using a 5-point scale: 0 – no signals, 1 – minimal occasional signals, 2 – mild scattered signals, 3 – moderate scattered signals, and 4 – abundant signals.

### Immunological assays

Antibody responses against IAV-S nucleoprotein (NP) were measured by using a commercial blocking ELISA (IDEXX, Montpellier, France), following the manufacturer’s recommendation. Results are expressed as a sample to negative control ratio (S/N ratio). Samples with an S/N ratio below 0.6 were considered positive.

HA-specific IgG antibodies in plasma and HA-specific IgA and IgG in BALF samples were measured using indirect ELISAs as described previously (28). The plasma samples were initially diluted at 1:100 in the sample dilution buffer and a 10-fold-serial dilution was performed and tested. An arbitrary cutoff value equivalent to the mean plus five standard deviations of the optical density (OD) values of samples from the non-immunized control animals was calculated. End-point antibody titers were determined at the highest dilutions that had the OD value above the assay cutoff. Samples with undetectable antibodies at this first dilution (1:100) were considered negative and assigned a titer of 1:10 for graphical and statistical purposes. BALF samples were tested at a single dilution (1:5) and data are presented as the OD_405_ value of the samples.

Hemagglutination inhibition (HI) assay and interferon-gamma (IFN-γ) ELISpot were performed as previously described (27).

### Flow cytometry

Cryopreserved PBMCs were thawed and seeded in a round bottom 96-well culture plate at the density of 1 million live cells in 100 μL of complete RPMI-1640 medium (cRPMI) as described previously (29). The cells were stimulated with the H3N2 TX98 virus at an MOI of 2. A separate set of PBMCs were treated with a cocktail containing phorbol 12-myristate 13-acetate (PMA, 10 ng/mL) and ionomycin (1 μg/mL) to serve as a positive control, or with cRPMI only to serve as a negative control. At 12 hr post-stimulation, 100 μL of cRPMI containing 1 μg/mL of GolgiPlug (containing Brefeldin A, BD Biosciences, San Jose, CA) was added to the cells to block the intracellular transport process. At 18 hr post-stimulation, the cells were washed twice with PBS (pH 7.4). Live-dead staining was performed using Zombie NIR^TM^ dye (BioLegend, San Diego, CA), and the Fc receptor was blocked with an anti-Fc receptor antibody (BioLegend). The cells were stained with a cocktail of antibodies against three surface makers: CD3ε, CD4α, and CD8α. After being fixed and permeabilized with the Cytofix/Cytoperm solution (BD Biosciences), the cells were stained with a cocktail of antibodies against three intracellular cytokines: IFN-γ, TNF-α, and perforin. After three washes with Perm/Wash buffer, the cells were resuspended in 300 μL of FACS buffer and analyzed by using the CytoFlex cytometer (Beckman Coulter, Fremont, CA). Approximately 100,000 events were acquired for each sample, and data were analyzed using FlowJo software (BD Biosciences).

### Quantification of Viral Load

RNA was extracted from nasal swabs and BALF samples using the Quick RNA viral Kit (Zymo Research, Costa Mesa, CA) and the viral genomic copies were quantified using a real-time reverse transcription PCR (RT-PCR) (VetMax-Gold SIV Detection Kit, Life Technologies, Austin, TX). A standard curve was established using a chemically synthesized RNA fragment with known copy numbers, based on which the absolute copy numbers of viral RNA in each sample were estimated (27). The viral loads were reported as log_10_ copies per 100 μL of samples. Samples with a cycle threshold value above 38 were considered negative and assigned a value of 0.8, equivalent to the assay limit of detection.

### Statistical Analysis

The statistical analyses were performed using GraphPad Prism 9.0. The HI antibody titers were log_2_ transformed and analyzed using the mixed-effects model. Univariate data including lung consolidation score, lung microscopic lesion score, area-under-the-curve of the viral genome copies in nasal swabs, virus titers in BALF samples, and frequencies of T cells expressing cytokines, were analyzed by analysis of variance (ANOVA), followed by Tukey’s multiple comparison test.

## Institutional Review Board Statement

The animal experiments were approved by the University of Nebraska-Lincoln (UNL) Institutional Animal Care and Use Committee under protocol number 2228, which was approved on April 12, 2022.

## Data Availability Statement

Not applicable.

## References

1. Vincent AL, Lager KM, Anderson TK. 2014. A brief introduction to influenza A virus in swine. Methods Mol Biol 1161:243–58.

2. Anderson TK, Macken CA, Lewis NS, Scheuermann RH, Van Reeth K, Brown IH, Swenson SL, Simon G, Saito T, Berhane Y, Ciacci-Zanella J, Pereda A, Davis CT, Donis RO, Webby RJ, Vincent AL. 2016. A Phylogeny-Based Global Nomenclature System and Automated Annotation Tool for H1 Hemagglutinin Genes from Swine Influenza A Viruses. mSphere 1.

3. Anderson TK, Chang J, Arendsee ZW, Venkatesh D, Souza CK, Kimble JB, Lewis NS, Davis CT, Vincent AL. 2021. Swine Influenza A Viruses and the Tangled Relationship with Humans. Cold Spring Harb Perspect Med 11.

4. Sandbulte MR, Spickler AR, Zaabel PK, Roth JA. 2015. Optimal Use of Vaccines for Control of Influenza A Virus in Swine. Vaccines (Basel) 3:22–73.

5. Venkatesh D, Anderson TK, Kimble JB, Chang J, Lopes S, Souza CK, Pekosz A, Shaw-Saliba K, Rothman RE, Chen KF, Lewis NS, Vincent Baker AL. 2022. Antigenic Characterization and Pandemic Risk Assessment of North American H1 Influenza A Viruses Circulating in Swine. Microbiol Spectr 10:e0178122.

6. Neveau MN, Zeller MA, Kaplan BS, Souza CK, Gauger PC, Vincent AL, Anderson TK. 2022. Genetic and Antigenic Characterization of an Expanding H3 Influenza A Virus Clade in U.S. Swine Visualized by Nextstrain. mSphere 7:e0099421.

7. Karlsson I, Borggren M, Rosenstierne MW, Trebbien R, Williams JA, Vidal E, Vergara-Alert J, Foz DS, Darji A, Sistere-Oro M, Segales J, Nielsen J, Fomsgaard A. 2018. Protective effect of a polyvalent influenza DNA vaccine in pigs. Vet Immunol Immunopathol 195:25–32.

8. Borggren M, Nielsen J, Karlsson I, Dalgaard TS, Trebbien R, Williams JA, Fomsgaard A. 2016. A polyvalent influenza DNA vaccine applied by needle-free intradermal delivery induces cross-reactive humoral and cellular immune responses in pigs. Vaccine 34:3634–40.

9. Bragstad K, Vinner L, Hansen MS, Nielsen J, Fomsgaard A. 2013. A polyvalent influenza A DNA vaccine induces heterologous immunity and protects pigs against pandemic A(H1N1)pdm09 virus infection. Vaccine 31:2281–8.

10. Gorres JP, Lager KM, Kong WP, Royals M, Todd JP, Vincent AL, Wei CJ, Loving CL, Zanella EL, Janke B, Kehrli ME, Jr., Nabel GJ, Rao SS. 2011. DNA vaccination elicits protective immune responses against pandemic and classic swine influenza viruses in pigs. Clin Vaccine Immunol 18:1987–95.

11. Macklin MD, McCabe D, McGregor MW, Neumann V, Meyer T, Callan R, Hinshaw VS, Swain WF. 1998. Immunization of pigs with a particle-mediated DNA vaccine to influenza A virus protects against challenge with homologous virus. J Virol 72:1491–6.

12. Larsen DL, Olsen CW. 2002. Effects of DNA dose, route of vaccination, and coadministration of porcine interleukin-6 DNA on results of DNA vaccination against influenza virus infection in pigs. Am J Vet Res 63:653–9.

13. Sistere-Oro M, Lopez-Serrano S, Veljkovic V, Pina-Pedrero S, Vergara-Alert J, Cordoba L, Perez-Maillo M, Pleguezuelos P, Vidal E, Segales J, Nielsen J, Fomsgaard A, Darji A. 2019. DNA vaccine based on conserved HA-peptides induces strong immune response and rapidly clears influenza virus infection from vaccinated pigs. PLoS One 14:e0222201.

14. Mashima R, Takada S. 2022. Lipid Nanoparticles: A Novel Gene Delivery Technique for Clinical Application. Curr Issues Mol Biol 44:5013–5027.

15. Corbett KS, Edwards DK, Leist SR, Abiona OM, Boyoglu-Barnum S, Gillespie RA, Himansu S, Schafer A, Ziwawo CT, DiPiazza AT, Dinnon KH, Elbashir SM, Shaw CA, Woods A, Fritch EJ, Martinez DR, Bock KW, Minai M, Nagata BM, Hutchinson GB, Wu K, Henry C, Bahl K, Garcia-Dominguez D, Ma L, Renzi I, Kong WP, Schmidt SD, Wang L, Zhang Y, Phung E, Chang LA, Loomis RJ, Altaras NE, Narayanan E, Metkar M, Presnyak V, Liu C, Louder MK, Shi W, Leung K, Yang ES, West A, Gully KL, Stevens LJ, Wang N, Wrapp D, Doria-Rose NA, Stewart-Jones G, Bennett H, et al. 2020. SARS-CoV-2 mRNA vaccine design enabled by prototype pathogen preparedness. Nature 586:567–571.

16. Hald Albertsen C, Kulkarni JA, Witzigmann D, Lind M, Petersson K, Simonsen JB. 2022. The role of lipid components in lipid nanoparticles for vaccines and gene therapy. Adv Drug Deliv Rev 188:114416.

17. Cheng X, Lee RJ. 2016. The role of helper lipids in lipid nanoparticles (LNPs) designed for oligonucleotide delivery. Adv Drug Deliv Rev 99:129–137.

18. Yanez Arteta M, Kjellman T, Bartesaghi S, Wallin S, Wu X, Kvist AJ, Dabkowska A, Szekely N, Radulescu A, Bergenholtz J, Lindfors L. 2018. Successful reprogramming of cellular protein production through mRNA delivered by functionalized lipid nanoparticles. Proc Natl Acad Sci U S A 115:E3351–E3360.

19. Yu B, Wang X, Zhou C, Teng L, Ren W, Yang Z, Shih CH, Wang T, Lee RJ, Tang S, Lee LJ. 2014. Insight into mechanisms of cellular uptake of lipid nanoparticles and intracellular release of small RNAs. Pharm Res 31:2685–95.

20. Gorres JP, Lager KM, Kong WP, Royals M, Todd JP, Vincent AL, Wei CJ, Loving CL, Zanella EL, Janke B, Kehrli ME, Nabel GJ, Rao SS. 2011. DNA Vaccination Elicits Protective Immune Responses against Pandemic and Classic Swine Influenza Viruses in Pigs. Clinical and Vaccine Immunology 18:1987–1995.

21. Liew FY, Li Y, Millott S. 1990. Tumor necrosis factor-alpha synergizes with IFN-gamma in mediating killing of Leishmania major through the induction of nitric oxide. J Immunol 145:4306–10.

22. Darrah PA, Patel DT, De Luca PM, Lindsay RW, Davey DF, Flynn BJ, Hoff ST, Andersen P, Reed SG, Morris SL, Roederer M, Seder RA. 2007. Multifunctional TH1 cells define a correlate of vaccine-mediated protection against Leishmania major. Nat Med 13:843–50.

23. Roces CB, Lou G, Jain N, Abraham S, Thomas A, Halbert GW, Perrie Y. 2020. Manufacturing Considerations for the Development of Lipid Nanoparticles Using Microfluidics. Pharmaceutics 12.

24. Sun HS, JH; Sillman S.; Steffen D.; Vu, HL. 2019. Design and characterization of a consensus hemagglutinin vaccine immunogen against H3 influenza A viruses of swine. Vet Microbiol In Press.

25. Halbur PG, Paul PS, Frey ML, Landgraf J, Eernisse K, Meng XJ, Lum MA, Andrews JJ, Rathje JA. 1995. Comparison of the pathogenicity of two US porcine reproductive and respiratory syndrome virus isolates with that of the Lelystad virus. Vet Pathol 32:648–60.

26. Gauger PC, Loving CL, Khurana S, Lorusso A, Perez DR, Kehrli ME, Jr., Roth JA, Golding H, Vincent AL. 2014. Live attenuated influenza A virus vaccine protects against A(H1N1)pdm09 heterologous challenge without vaccine associated enhanced respiratory disease. Virology 471–473:93-104.

27. Sun H, Sur JH, Sillman S, Steffen D, Vu HLX. 2019. Design and characterization of a consensus hemagglutinin vaccine immunogen against H3 influenza A viruses of swine. Vet Microbiol 239:108451.

28. Kumari S, Chaudhari J, Huang Q, Gauger P, De Almeida MN, Liang Y, Ly H, Vu HLX. 2022. Immunogenicity and Protective Efficacy of a Recombinant Pichinde Viral-Vectored Vaccine Expressing Influenza Virus Hemagglutinin Antigen in Pigs. Vaccines (Basel) 10.

29. Chaudhari J, Liew CS, Workman AM, Riethoven JM, Steffen D, Sillman S, Vu HLX. 2020. Host Transcriptional Response to Persistent Infection with a Live-Attenuated Porcine Reproductive and Respiratory Syndrome Virus Strain. Viruses 12.

